# MicrobeMod: A computational toolkit for identifying prokaryotic methylation and restriction-modification with nanopore sequencing

**DOI:** 10.1101/2023.11.13.566931

**Authors:** Alexander Crits-Christoph, Shinyoung Clair Kang, Henry H. Lee, Nili Ostrov

**Affiliations:** Cultivarium, 490 Arsenal Way, Watertown MA 02472, USA

## Abstract

Bacteria and archaea use restriction-modification (R-M) systems to distinguish self from foreign DNA by methylating their genomes with DNA methyltransferases with diverse sequence specificities, and these immunity systems often vary at the strain level. Identifying active methylation patterns and R-M systems can reveal barriers to the introduction of recombinant DNA or phage infection. Here, we present the computational MicrobeMod toolkit for identifying 5mC and 6mA methylation sequence motifs and R-M systems in bacterial genomes using nanopore sequencing of native DNA. We benchmark this approach on a set of reference *E. coli* strains expressing methyltransferases with known specificities. We then applied these analyses to 31 diverse bacterial and archaeal organisms to reveal the methylation patterns of strains with previously unexplored epigenetics, finding that prokaryotic 5-methylcytosine may be more common than previously reported. In summary, MicrobeMod can rapidly reveal new epigenetics within a prokaryotic genome sequenced with Oxford Nanopore R10.4.1 flow cells at sequencing depths as low as 10x and only requires native DNA. This toolkit can be used to advance fundamental knowledge of bacterial methylation and guide strategies to overcome R-M barriers of genetic tractability in non-model microbes.

## Introduction

### Methylation for self recognition and defense

Bacteria and archaea possess a myriad of defense systems that identify and protect against exogenous phage, plasmids, and mobile elements^1^. The most ubiquitous of these defense systems within the bacterial domain is restriction-modification (R-M)^2^, which was also the first type to be identified^3^. Prokaryotic R-M systems use DNA methyltransferases to methylate endogenous DNA at specific sequence motifs, allowing cognate restriction enzymes targeting those sequences to cleave foreign unmethylated DNA, and not methylated endogenous DNA. This method of defense is widespread with 80% of prokaryotic genomes and 10% of plasmids^4^ encoding for at least one R-M system^1,5^. Some species such as *Helicobacter pylori* can have over 15 unique sequence motifs methylated in their genomes^6^.

Three forms of DNA methylation are thought to be common in prokaryotes: 6-methyladenosine (6mA), 4-methylcytosine (4mC), and 5-methylcytosine (5mC)^7^. The dominant form of methylation involved in R-M is thought to be 6mA, with one study finding it comprising 75% of methylation motifs across a set of 230 bacterial species^8^. However, not all prokaryotic methylation is involved in R-M, as methylation is also known to play roles in cellular regulation which can be mediated by “orphan” methyltransferases without cognate restriction enzymes like the Dam and Dcm methylases of *Escherichia coli*^9^. Previous studies have estimated that DNA methylation of any form exists in over 90% of prokaryotes^8^.

Across diverse microorganisms, restriction modification has been identified as a frequent barrier for genetic tractability^10–19^. This obstacle is expected, as the natural role of R-M is to identify and destroy foreign DNA. The degree to which R-M impacts transformation efficiency varies by strain and may not present a barrier for genetic tractability in all strains. However, circumventing native R-M systems has been shown to increase transformation efficiency by several orders of magnitude in many strains^20–22^ which can make the difference between successful and unsuccessful transformations^23^. Multiple experimental methods for circumventing R-M have been demonstrated, which include switching the methylation status of the donor strain^23,24^, incubating plasmids in cell lysate with inhibited restriction enzymes^25^, heterologous^11,20^ or cell-free^26^ expression and incubation with native methyltransferases, and engineering of plasmids to remove any sequences matching the R-M motif specificities of a given strain^27^. However, most of these approaches have an *a priori* requirement of identifying the methylation status, methylation sequence motifs, and/or R-M genes for a given target strain.

### Sequencing methods to detect methylation patterns and restriction-modification

In the last decade, approaches have been developed to identify methylated bases in genomes using both single molecule, real-time (SMRT) sequencing from Pacific Biosciences (PacBio)^28^, and more recently, nanopore sequencing from Oxford Nanopore^7,29,30^. The REBASE database is the most comprehensive source of R-M systems, providing methylation sequence motifs and restriction modification systems for over 4890 strains with links to publicly available SMRT sequencing data^31^.

Due to the rapid maturation and accessibility of Oxford Nanopore sequencing there is a strong interest to develop methods to detect modified nucleotides and methylation patterns using this platform. For example, the software Nanodisco^32^, developed for the Oxford Nanopore R9.4 series of flow cells, enabled high-throughput analyses of methylation sequence motifs in microbial genomes^33^. To identify methylated bases, this approach compares the electrical current difference detected by the nanopore between native DNA with intact methylation and whole-genome amplified DNA without methylation. Methylation motifs are then identified by passing modified genomic sites to the MEME suite^34^ for motif discovery. The software Snapper also compares native DNA to a whole genome amplified control, with the additional implementation of a greedy motif selection algorithm for improved motif identification^35^. Both methods were developed for R9.4 flow cells and required both native and amplified DNA to obtain methylation information for each strain.

To identify active enzymes in R-M systems, these genes must first be annotated in a strain’s genome. Multiple computational tools exist for the annotation of defense system genes in prokaryotic genomes. These approaches are homology-based and search genomes using sets of profile-HMMs targeted to the methyltransferase and restriction enzyme classes found in each R-M system type^4^. DefenseFinder and PADLOC systematically identify operons homologous to a broad array of known defense systems, including R-M systems^1,36^. DNA Methylase Finder specifically annotates all potential DNA methyltransferases in a genome and further distinguishes DNA methyltransferases from methyltransferases with other biological activities ^33^.

The rapid rate of technology development around long-read sequencing presents both opportunities and challenges. Oxford Nanopore released the R10.4.1 flow cell model during 2022, and multiple studies have reported significant improvements to read quality and consensus assembly accuracy in combination with the corresponding updated chemistry (V14)^37,38^. Oxford Nanopore further released modified base calling models for both 5mC and 6mA methylation that were compatible with R10.4.1 flow cells. However, these chemistry and base calling changes were incompatible with software packages such as Tombo and Nanopolish^39^, that existing methylation motif discovery tools such as Nanodisco and Snapper depended upon.

We have developed MicrobeMod as a singular workflow that integrates disparate packages to identify and analyze prokaryotic methylation nanopore sequencing and that is compatible with R10.4.1 flow cells. Crucially, this pipeline only requires sequencing reads from native DNA as input by using the modified base calling models in the Dorado basecaller. This approach is able to successfully recover methylated sites and their corresponding methylation motifs even with sequencing coverages as low as 10x. MicrobeMod contains simple and fast workflows for both the identification of methylation motifs in nanopore data and the annotation of R-M genes and operons, with the goal of enabling biological analyses that link gene predictions to observed active methylation patterns within a genome.

## Results

### Implementation overview

The MicrobeMod toolkit was implemented in Python 3.9 and has two main commands that correspond to two independent and complementary pipelines, *MicrobeMod call_methylation* and *MicrobeMod annotate_rm* (**Figure 1**). The *Call_methylation* pipeline is used to identify methylated sequence motifs in the genome. The input for the *call_methylation* pipeline is a BAM file of reads with assigned methylation probabilities output by the Dorado basecaller mapped to a genome assembly, and a FASTA file of that corresponding assembly. The *annotate_rm* pipeline is used to identify R-M methyltransferase and restriction enzyme genes identified in the genome, and group them by operonic context. The input for the *annotate_rm* pipeline is a FASTA genome assembly of any origin. Below we provide a description of each pipeline as well as examples for its implementation, and for more details, refer to *Materials and Methods*.

**Figure 1.**
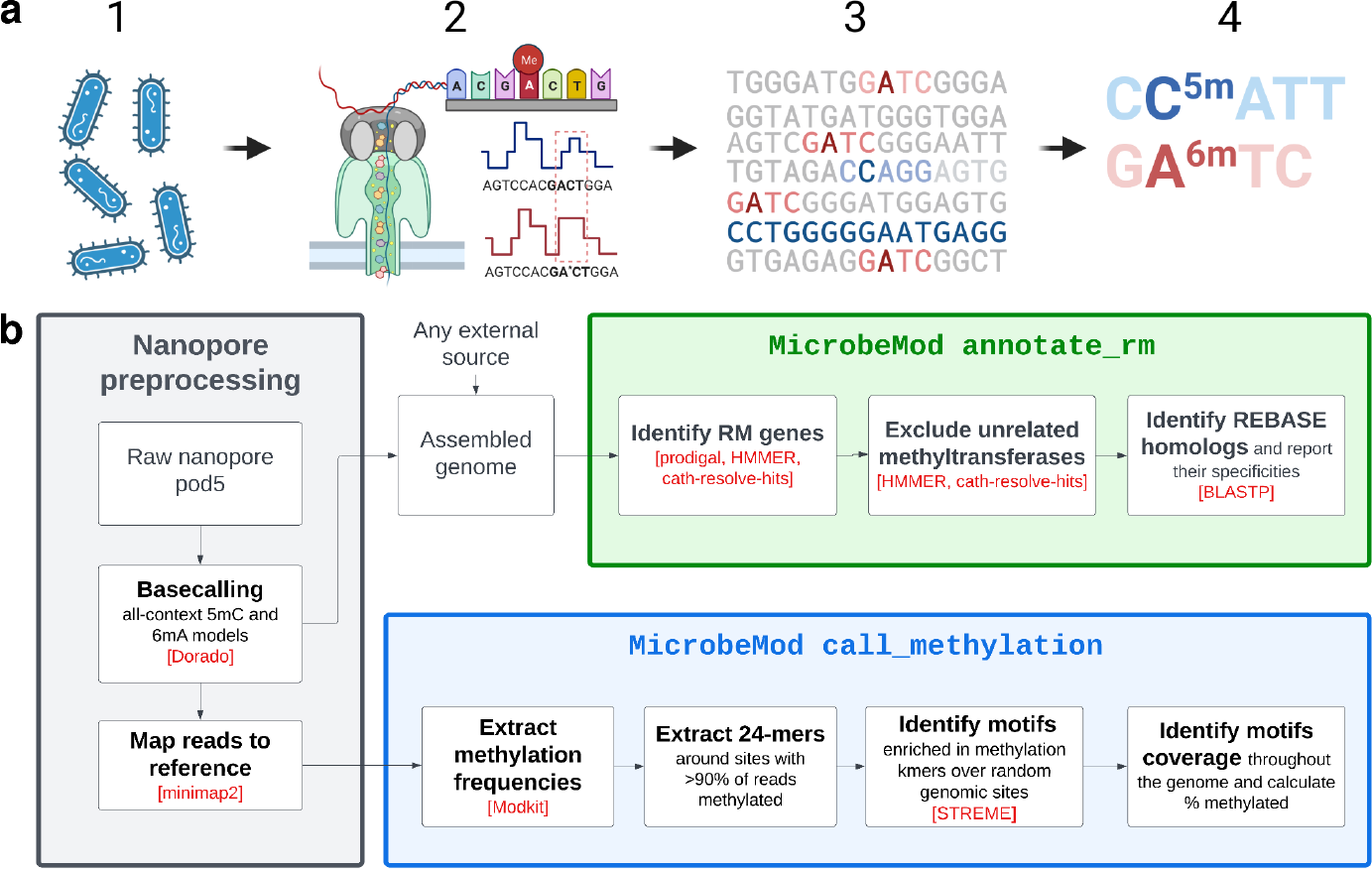
Overview of the MicrobeMod toolkit. **(a)** Conceptual overview of identification of methylated motifs in prokaryotic genomes. Native microbial DNA (1) is extracted and sequenced via nanopore sequencing (2), which detects differences in the current signal associated with modified bases. Sequencing reads are then mapped to genomes (3). MicrobeMod can be used to identify sequence motifs corresponding to the specificities of active methyltransferases in the strain (4). **(b)** Technical workflow for the MicrobeMod toolkit. Preprocessing steps are shown in gray, while the steps shown in blue and green correspond to the pipelines run by the *MicrobeMod call_methylation* and *MicrobeMod annotate_rm* commands, respectively. External dependencies are labeled in red. Created with BioRender.com.

The *MicrobeMod call_methylation* pipeline first calls Modkit to identify methylated sites and extract the methylation frequencies of reads at those sites, filtering sites based on both a minimum required coverage and percent of mapped reads methylated. By default, the 24 base pairs around positions for which at least 90% of reads are methylated are extracted and passed to STREME^40^ to identify significantly enriched sequence motifs compared to the genomic background. Resulting motifs are filtered by MicrobeMod with an e-value cutoff of 0.1, all genomic occurrences of the motifs are identified, and the fraction of those occurrences that contain a methylated site is reported as the “methylation coverage”. Finally, the top two most common positions that are methylated within a motif, and the frequency that each is methylated, are compared to identify the exact methylated bases within an identified motif. The final output is a table containing information for all of the methylation motifs identified in a genome, including their methylation types, frequencies, and methylated positions (**Table 2**)

The *MicrobeMod annotate_rm* pipeline annotates genes and operons involved in R-M in a provided genome by calling genes with prodigal^41^ and then searching for methyltransferase and restriction enzyme genes using HMMER^42^. R-M systems typically include a restriction enzyme and methyltransferase in an operonic context with each other. To identify operonic R-M systems, we assigned genes to operons based on a 10 gene window. The output of MicrobeMod *annotate_rm* preferentially ranks complete R-M operons over singleton genes (**Table 3**). Type IV restriction enzymes, which target methylated and modified DNA and thus are not associated with any corresponding methyltransferase, are considered operonic alone.

To provide information on the methylation specificity of identified R-M genes, the final step of MicrobeMod is to compare all identified genes to a set of characterized enzymes from the REBASE database via BLASTP. The top REBASE homolog hit for each gene in the user’s provided genome is reported, and in cases where the protein identity of the homolog is >75%, the methylation type and specificity of the homolog are reported, as methylation type, and to a lesser degree motif specificity, can sometimes be predicted via homology. The final output is a table of all potential genes identified as involved in R-M within the provided genome, ranked by their operonic context, and including all subtype and homolog information.

### Experimental validation of the MicrobeMod pipeline

The accuracy of Dorado-based methylation calling using Oxford Nanopore R10.4.1 flow cells and the MicrobeMod pipeline was tested on a set of *E. coli* strains with known methylation patterns. Mean sequencing depths for these strains ranged from 47x to 304x (**Table 1**), with a mean read length of 4166 bp. First, as a negative control, we sequenced a methylation-deficient *E. coli* strain (NEB C2925H). MicrobeMod correctly identified an absence of any significant methylation motifs in this strain (**Figure 2a**). Very few sites were identified as methylated total: using a default 66% cutoff for percent of reads methylated at a site, only 661 6mA sites were called, corresponding to a genome-wide methylation frequency <0.014%; with a cutoff of 90%, only 13 6mA sites were identified (0.00028%). We noticed that these 13 false positive sites were flanked by homopolymer stretches of C and G which may be responsible for abnormal current signals that could result in methylation miscalls. In MicrobeMod, 66% read methylation at a site is required to identify a site as methylated, while 90% is required to further include that site in motif calling (each percentage is tunable via their respective parameters). Zero 5mC sites were identified in this strain at either cutoff.

**Figure 2.**
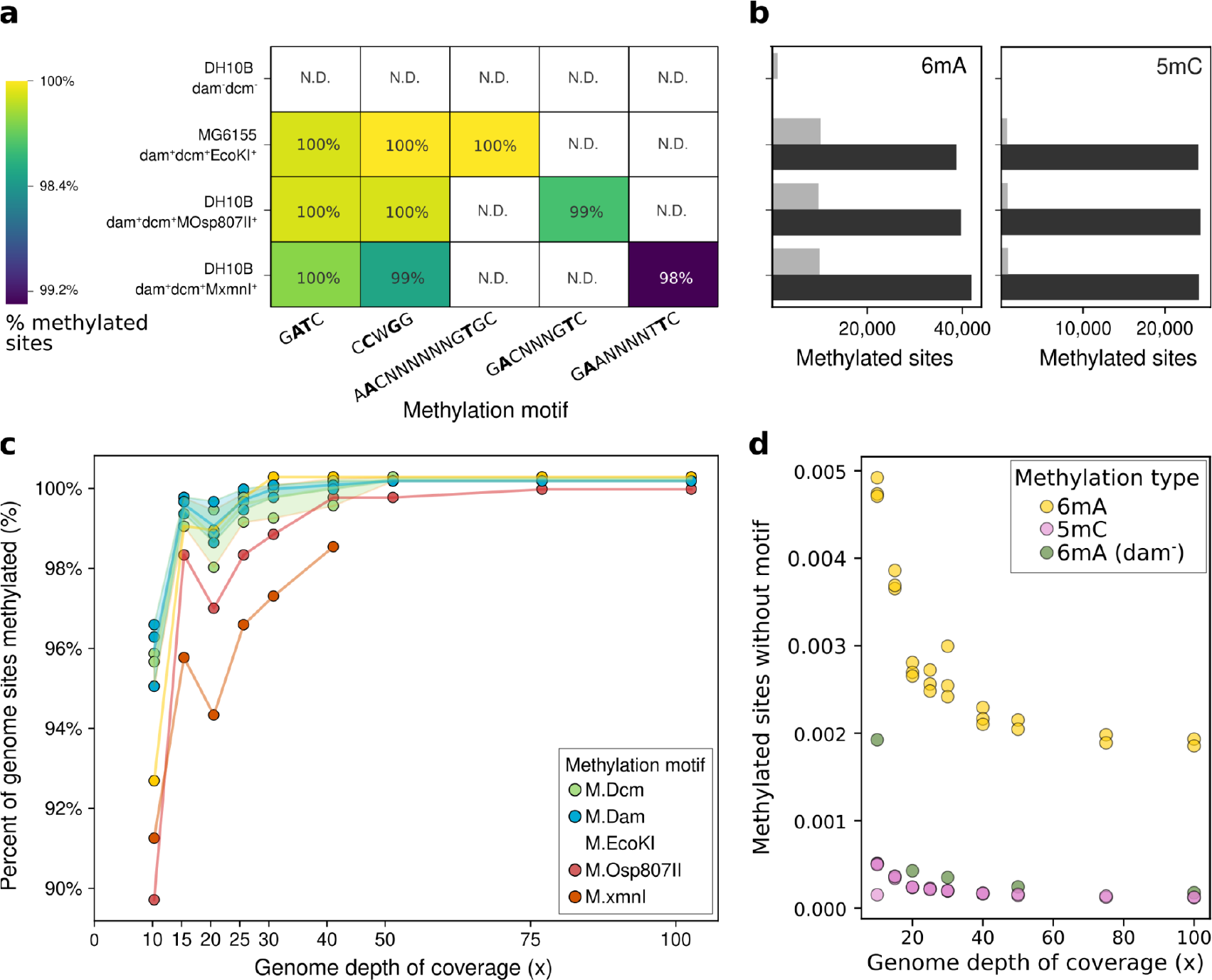
Benchmarking MicrobeMod using *E. coli* strains with known methyltransferase specificities. **(a)** Methylation motifs identified by MicrobeMod on four benchmarking *E. coli* strains with known methylation patterns. The percent of all sites in the genome matching the identified methylated motif is shown. N.D. indicates the motif was not detected. **(b)** The total number of 6mA (left) and 5mC (right) methylated bases in the genome of each strain in (a), as output by Modkit and filtered by MicrobeMod. Black bars indicate motif-associated sites. Gray bars indicate sites with no associated motif. **(c)** Downsampling true-positive benchmark of MicrobeMod. Each benchmark strain was downsampled to a normalized mean genome depth of coverage (x-axis) and the percentage of motif sites in the genome correctly identified as methylated (y-axis). **(d)** Downsampling false-positive benchmark of MicrobeMod, where the numbers of sites not matching any known motif were assessed (y-axis).

**Table 1:**
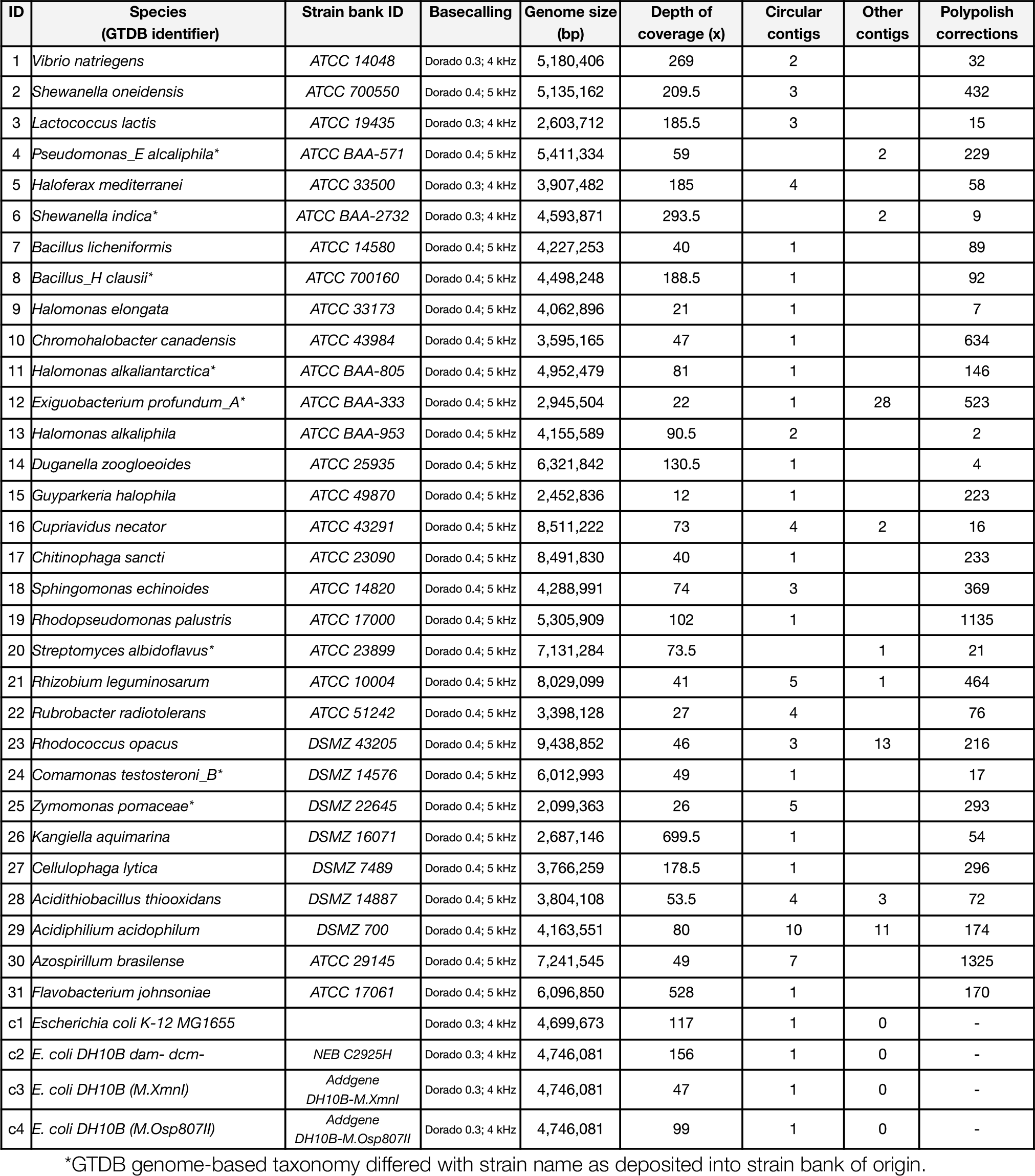
Strains included in this study.

**Table 2:** Example MicrobeMod call_methylation output, for *E. coli MG1655*.

| Motif | Motif raw | Methylation type | Genome sites | Methylated sites | Methylation coverage | Average Percent Methylation per site | Methylated position 1 | Methylated position 1 percent | Methylated position 2 | Methylated position 2 percent |
| --- | --- | --- | --- | --- | --- | --- | --- | --- | --- | --- |
| GATC | 1-YGNYG<br>ATCNBNH<br>NVN | a | 38248 | 38216 | 0.999 | 0.93 | 2 | 50 | 3 | 50 |
| CCWGG | 2-NNSVR<br>CCWGGY<br>BSNN | a | 24100 | 14242 | 0.591 | 0.83 | 3 | 100 | NA | 0 |
| GCACNN<br>NNNNGTT | 3-GCACB<br>VNVNNG<br>TTN | a | 595 | 595 | 1 | 0.9 | 3 | 48.595 | 12 | 48.099 |
| No Motif Assigned | NA | a | NA | 9163 | NA | 0.77 | NA | NA | NA | NA |
| CCWGG | 1-NNNNR<br>CCWGGY<br>NNNN | m | 24100 | 24086 | 0.999 | 0.94 | 2 | 48.87 | 4 | 48.87 |
| No Motif Assigned | NA | m | NA | 542 | NA | 0.72 | NA | NA | NA | NA |

**Table 3:**
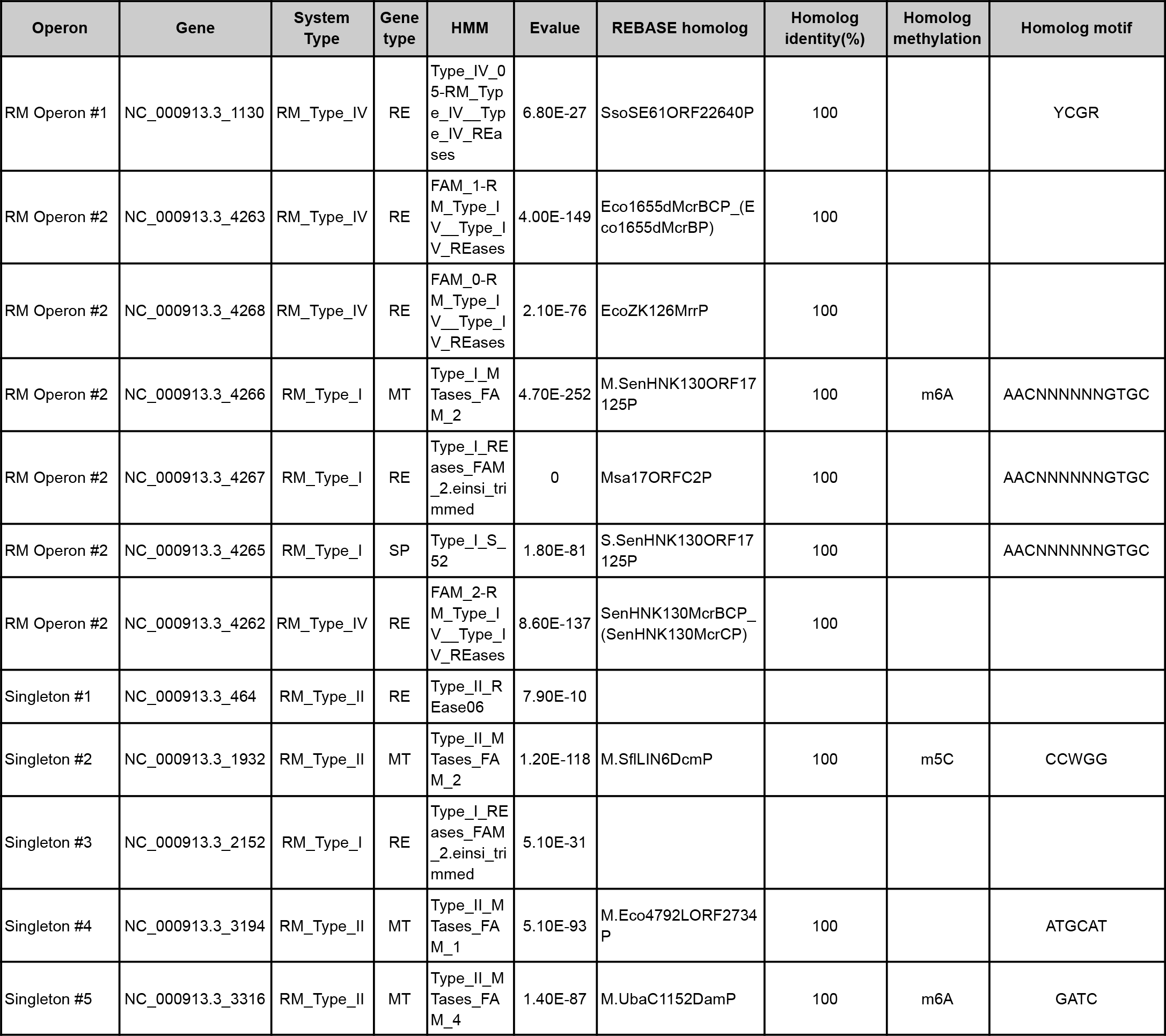
Example MicrobeMod annotate_rm output, for *E. coli MG1655*.

We next sequenced *E. coli* MG1655, with active Dam methyltransferase (which methylates 6mA in motif G**AT**C), Dcm methyltransferase (which methylates 5mC in motif C**C**W**G**G; W=A or T), and the EcoKI (which methylates 6mA in motif A**A**CNNNNNNNG**T**GC) R-M system. MicrobeMod identified the exact canonical motifs for all three methyltransferases and reported approximately 100% of all sites with this sequence motif as methylated (**Figure 2a**). One potential confounding issue observed was a significant fraction of CC**W**GG sites miscalled with 6mA at the 3rd position (called as methylated in 59% of motif occurrences). This presumably inaccurate modified base calling behavior only seemed to occur in Dcm+ *E. coli* strains, and may be due to adjacent modified bases altering the current signal in a way that resembles a 6mA modification to the base calling model. However, the far greater assignment of CCWGG to 5mC versus 6mA (100% vs 59%, respectively) indirectly infers the correct methylation status of this motif, and we did not observe this behavior elsewhere in the data.

To further test the accuracy on more unusual motifs, we sequenced two *E. coli* DH10B strains expressing the heterologous methyltransferases M.Osp807II (which methylates 6mA in motif G**A**CNNNG**T**C) and M.xmnI (which methylates 6mA in motif G**A**ANNNNT**T**C)^*43*^. These methyltransferases are originally from *Olsenella sp. oral taxon 807* (phylum Actinomycetota) and *Xanthomonas phaseoli pv. Manihotis* (phylum Pseudomonadota)^43^. In each case, MicrobeMod recovered the exact known bipartite motifs of these methyltransferases, identified >98% of genomic sites methylated, and reported no false positive motifs (**Figure 2a**). In each case these motifs were identified in a background genome also expressing Dam and Dcm methyltransferases. Together, these results provided confidence that Dorado-base calling with R10.4.1 flow cells can accurately identify both 5mC and 6mA methylated sites, and MicrobeMod can identify palindromic and non-palindromic motifs, bipartite motifs, multiple motifs within the same genome, and motifs associated with non-model methyltransferases.

The success and accuracy of consensus methylation calling can be dependent upon the depth of coverage of the sequenced genome. To determine the coverage required for reliable methylation calling, we sub-sampled each of the benchmarking genomes at set coverage thresholds starting with a maximum of 100x coverage down to a minimum of 10x coverage, re-running MicrobeMod on each subsampled set of reads. In all sub-sampled datasets, MicrobeMod was able to identify the correct methylation status of each benchmark strain (**Figure 2c**). Furthermore, we found that the percentage of genomic sites of the expected sequence motifs were identified as methylated in over 90% of cases, even with coverage as low as 10x. This result indicated that MicrobeMod is able to correctly recover methylation motifs in genomes that are sequenced as low as 10x coverage.

Methylation can also occur at sites not associated with motifs that could represent either instances of off-target methylation by the methyltransferases or technical false positives. We evaluated whether low coverage increased the rate of methylated sites not associated with motifs. Overall, false positive or off-target methylated sites were only slightly more common at coverages as low as 10x than they were at 100x (**Figure 2d**). In particular, false positive / off-target methylated sites were at or near 0 for 5mC. In the three strains with at least one active methyltransferase, 6mA false positive / off-target sites were ∼2.5 times as common at 10x as they were at 100x depth of coverage (**Figure 2b,2d**). However, these sites did not impact the motif calling results, indicating they had little systematic impact. Intriguingly, the false positive rate of the methylation deficient *E. coli* strain was substantially lower (**Figure 2d**). This could potentially indicate that some methylated sites not associated with motifs in *E. coli MG1655* could be due to off-target biological activity of the methyltransferases, as opposed to being technical false positives. In combination, these results indicate that MicrobeMod was capable of accurately identifying methylation motifs in genomes largely independent of the depth of coverage and at coverages as low as 10x sequencing depth.

### Analyzing diverse prokaryotic epigenomes using MicrobeMod

We next sequenced native DNA from a diverse collection of 30 bacterial strains and one archaeal strain and analyzed these data with MicrobeMod to infer their methylation status. Selected organisms included strains from phyla Bacteroidota, Bacillota, Pseudomonadota, Actinomycetota, and Euryarchaeota with genomes ranging from 2-9.4 Mb in size. Nanopore sequencing coverage of each strain ranged from 12x to 699x, with a mean coverage of 40x, and an average read length of 4000 bp (**Table 1**). Genomes were assembled *de novo* from long-reads with Flye, which resulted in highly contiguous assemblies: 22 of these genomes were circularized and 15 of them represented a significant improvement upon the contiguity of the closest reference genome in NCBI. To test the consensus genome accuracy of nanopore-alone versus a hybrid assembly approach, PolyPolish was run on each genome using additional Illumina sequencing reads: on average, there were only 239 PolyPolish corrected errors across each genome, consistent with Q50+ consensus accuracies of nanopore-only genomes reported in other recent benchmarks^52^.

Assembled genomes and corresponding read mappings were next used as input for MicrobeMod to infer methylation status. We first analyzed 9 strains which had previously published methylation data based on PacBio sequencing and made available through the online database REBASE. Of the 19 6mA methylation motifs identified by MicrobeMod across these 9 strains, 15 had an exact corresponding match as reported in REBASE, and another 4 motifs with a “close” match (**Figure 3a**). Close matches included cases where the motif differed by 1 or 2 base pairs or was otherwise overly specific: for example, for *Shewanella oneidensis MR-1* REBASE reports an **ATCGAT** motif while MicrobeMod identified a more specific **ATCGAT**NRNS motif.

**Figure 3:**
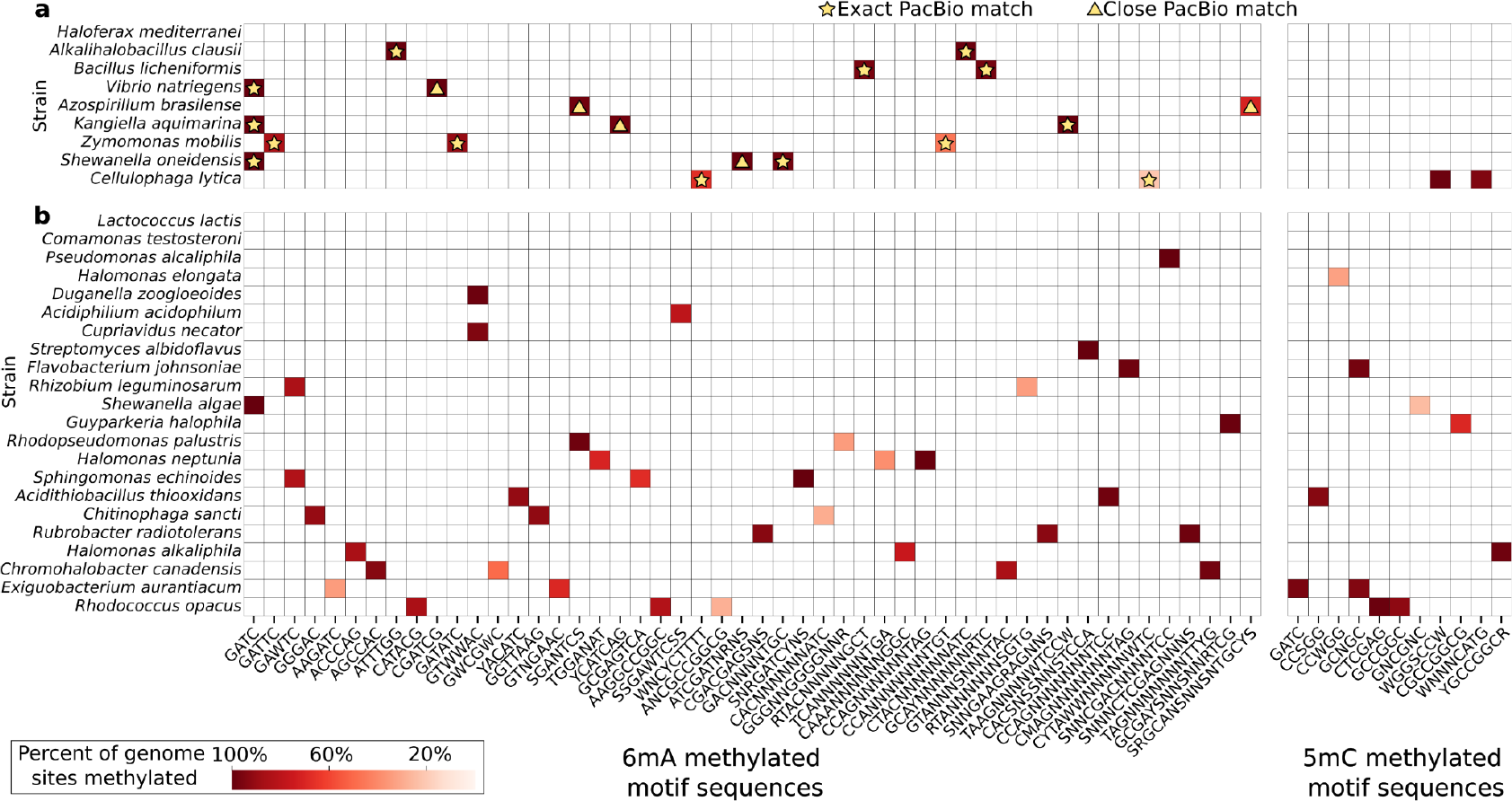
Methylation patterns detected by MicrobeMod in diverse prokaryotic strains. **(a)** Methylation motifs in 9 strains for which data is available in REBASE. Color gradient indicates percent of methylated motif in the respective genome. White color indicates the motif was not found. A star indicates the exact motif was identified by both REBASE and MicrobeMod, and very similar motifs are marked by a yellow triangle. Motifs follow the IUPAC code for ambiguous nucleotides. **(b)** Methylation motifs in 21 strains for which no data is available in REBASE. Color scheme is the same as panel a.

Overall, these results indicate a high degree of congruence between PacBio- and Oxford Nanopore-based methylation calling approaches. Most of the strains with REBASE information had no 5mC methylation reported either by MicrobeMod or REBASE. However, *Cellulophaga lytica* had two examples of 5mC methylation motifs reported by MicrobeMod not reported by REBASE: WG**G**S**C**CW and WNNN**C**AT**G**, which had a robust genomic methylation coverage of 98.7% and 94.9% respectively. These calls are likely correct, as this strain also encodes for homologs of the methyltransferases M.Cly7489ORF997P and M.Cly7489ORF1631P with reported specificities of G**G**N**C**C and **C**AT**G**; it is possible that these methylation motifs are missed by the PacBio approach due to potentially lower specificity for detection of 5mC^7^. Finally, one strain in this set (*Haloferax mediterranei*) had no methylation detected by MicrobeMod, but has 4mC methylation with two associated sequence motifs^44^. This result confirms that the currently available all-context 5mC modified base calling models used in this study are specific to 5mC and there is no indication of misclassification of 4mC.

We further analyzed the full set of 31 sequenced prokaryotic strains using MicrobeMod, including 22 bacterial strains without any previously reported epigenetic data (**Figure 3b**). Of these, we identified a mean of 2.1 methylation motifs per strain, and 93% of the strains had any detectable methylation motifs containing 5mC or 6mA. Notably, we did not detect any methylation motifs for *Lactococcus lactis* and *Comamonas testosteroni*. The 6mA methylation was considerably more common than 5mC, with 87% of strains having at least one 6mA motif and 29% of strains having at least one detectable 5mC motif. This prevalence of 5mC methylation was significantly higher than previously reported based on PacBio sequencing of a large cohort of diverse strains^8^. The most complex methylome detected was *Rhodococcus opacus*, with five active methylation motifs, followed by *Exiguobacterium aurantiacum, Chromohalobacter canadensis*, and *Cellulophaga lytica*, each with four methylation motifs. The G**AT**C motif associated with the orphan Dam methyltransferase from *E. coli* was also identified in four members of the Gammaproteobacteria, consistent with its widespread function in gene regulation in this clade^45^. Correspondingly, the G**A**W**T**C motif, associated with homologs of the orphan CcrM methyltransferase in the alphaproteobacterium *Caulobacter crescentus* was detected in two of the Alphaproteobacteria in the set (*Rhizobium leguminosarum* and *Sphingomonas echinoides*). This was also consistent with the widespread function of this methylase in gene regulation among the Alphaproteobacteria^46^.

We next annotated R-M genes within this collection of microbial genomes using the *MicrobeMod annotate_rm* pipeline. The mean number of R-M operons detected per genome was 1.7, and across all 31 microbes 26 Type I, 19 Type II, 16 Type IIG, 3 Type III, and 13 Type IV R-M systems were identified (**Figure 4a**). Type II methyltransferases were the most common of all singleton methyltransferase or restriction enzyme genes not found in an operonic context with other R-M genes, consistent with the known high frequencies of orphan Type II methyltransferases (**Figure 4b**). Many of these orphan Type II methyltransferases are likely inactive, such as the M.Eco1655ORF6040P ORF in *E. coli* MG1655, or these genes may play more unknown roles beyond DNA methylation.

**Figure 4:**
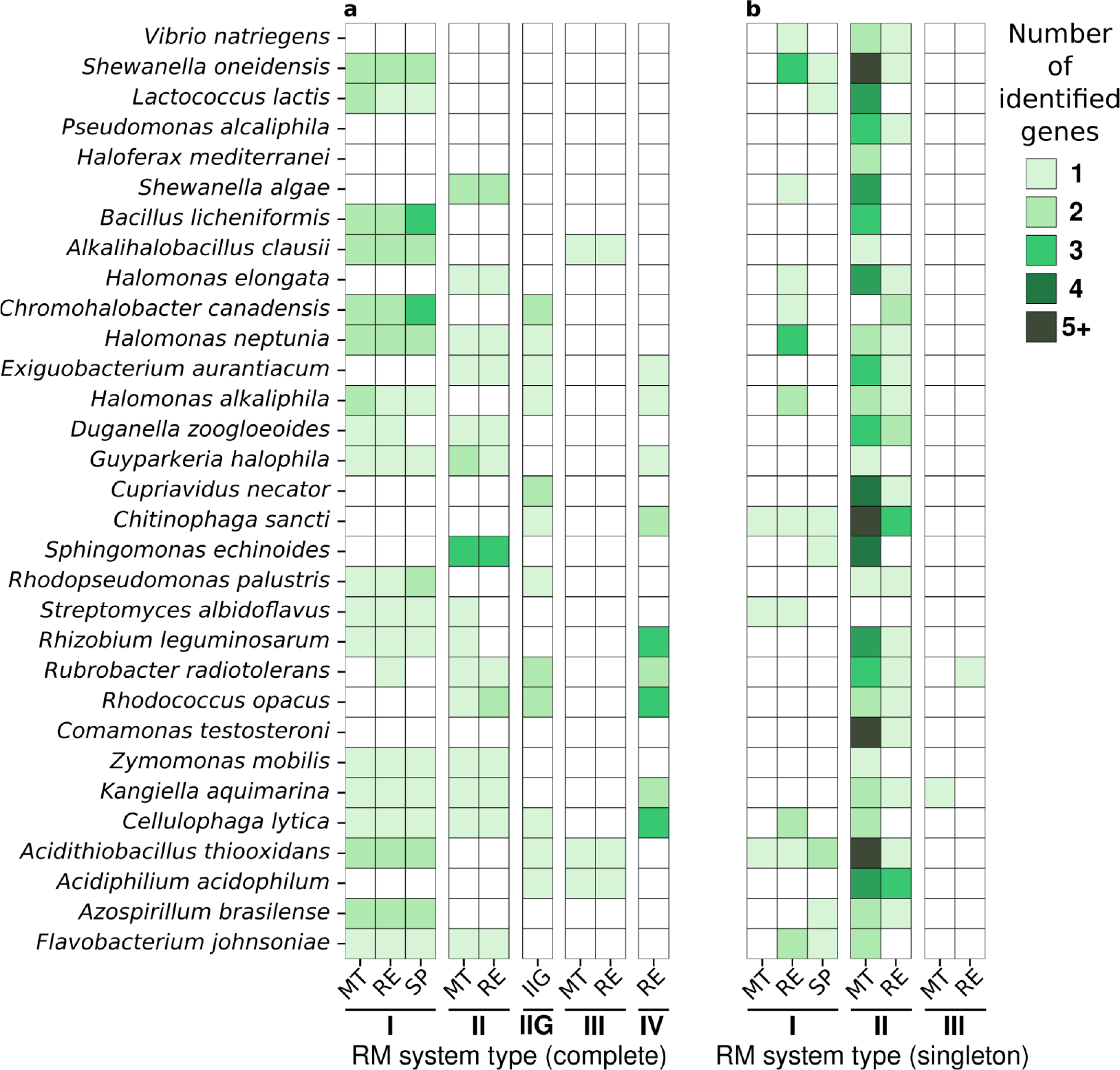
MicrobeMod annotation of restriction-modification genes in diverse prokaryotic strains. Each row represents a strain annotated using *MicrobeMod annotate_rm*. The number of genes identified under each restriction-modification (R-M) type is represented by the color gradient. **(a)** “Complete” R-M systems represent instances where all required subunits of a specific R-M type were detected within 10 genes of each other. **(b)** “Singleton” R-M genes are genes that are not associated with a complete R-M system. Abbreviations: MT (methyltransferase), RE (restriction enzyme), SP (specificity subunit), IIG (Type IIG restriction enzyme which also includes a methyltransferase domain).

Ideally, observed methylation motifs could be directly associated with their respective R-M systems, but inferring these relationships may be challenging in practice. While MicrobeMod identifies both active methylation motifs and R-M genes, it does not directly assign each methylation motif to its corresponding R-M genes. In many cases, users can link gene annotations directly to observed active methylation patterns using the information provided by the MicrobeMod pipelines. Specifically, R-M genes and observed motifs can be linked by considering the R-M system types, and the closest homolog with its reported methylation motif.

For example, in the halophile *Halomonas alkaliphila, MicrobeMod annotate_rm* identifies a Type I R-M system, a Type IIG R-M system, and two orphan Type II methyltransferase genes. Correspondingly, *MicrobeMod call_methylation* identifies three methylated motifs: CAA**A**NNNNNNGGC, ACCC**A**G, and YGC**CG**GCR. The Type I R-M system has 96% amino acid identity to the M1.HspMS1ORF6835P from *Halomonas species MS1*, known to have 6mA methylation, and therefore can be assigned to the bipartite CAA**A**NNNNNNGGC motif, consistent with Type I R-M methylation motifs. Correspondingly, the Type IIG has 79% amino acid identity to the HspXH26ORF9735P enzyme, which also has a 6mA specificity - and therefore likely can be assigned to the ACCC**A**G motif, which is short and non-palindromic, features of Type IIG methylation motifs. Finally, of the two orphan methyltransferases, one has 60% amino acid identity to M.Bgl949ORF22065P with known 5mC specificity and therefore likely can be assigned to the YGC**CG**GCR. In this manner, MicrobeMod output can be used to not just identify R-M genes and methylation motifs, but to specifically link both together.

## Discussion

In conclusion, the MicrobeMod pipeline is able to accurately identify active methylation motifs using nanopore sequencing and annotate their corresponding R-M genes and systems within genomes. It is a versatile pipeline that can aid in interpretation of bacterial methylome data and to inform methylation-aware transformation strategies in a synthetic biology context. By applying this approach to a diverse set of bacterial isolates with previously unreported methylation data, we identified a significantly higher prevalence of 5mC methylation in bacteria than previously reported by SMRT cell sequencing^8^. SMRT cell sequencing may be less sensitive to 5mC detection than nanopore sequencing, which could have resulted in a prior underestimate of the frequency of 5mC methylation in prokaryotic genomes^7^.

There are also some limitations to be considered with methylome analysis in MicrobeMod, and in general with nanopore-based approaches. As there is currently no all-context 4mC modified base calling model available, this method is incapable of detecting 4mC in prokaryotic genomes in its current state. One of our sequenced strains (*H. mediterranei*) should have active 4mC methylation^44^, and thus this data may be useful in testing or benchmarking 4mC detection in the future. Second, we observed some degree of methylation miscalling: specifically 6mA miscalls in the presence of the C**C**W**G**G 5mC motif. The *E. coli* strain data in this study was sequenced with a 4 kHz sampling rate (**Table 1**); the Oxford Nanopore switch to 5 kHz, and the corresponding updated base calling models, may have largely solved this issue, which we have not yet observed in the other strains sequenced with the newer 5 kHz sampling rate. More broadly, detecting methylation with nanopore sequencing requires building a machine learning model that can distinguish target nucleotide modifications from the full variety of possible DNA modifications observed across diverse organisms. Thus, a theoretical risk could remain when modified base calling models are challenged with alternative base modifications they have not seen prior or in their training sets.

Motif calling with MicrobeMod is sometimes imperfect. In particular, if complex motifs are rare throughout a given genome, there may be insufficient motif occurrences to properly identify the motif’s exact specificity, resulting in a called motif that is slightly more specific than the true specificity of the enzyme. Because MicrobeMod can be run on a given bacterial genome on a contemporary 8-core laptop in 10-20 minutes, it can be useful to perform the analyses with various levels of methylation confidence stringency and percent methylation cutoffs to observe any potential differences in the identified motif sequences. In our dataset, the most complex methylome of any strain had five unique sequence motifs identified. Performance of MicrobeMod has not been benchmarked on strains with more complex methylomes, such as *Helicobacter pylori* strains.

While Oxford Nanopore library chemistry and base calling models can be subject to rapid change, MicrobeMod’s input is a standardized BAM format irrespective of flow cell chemistry, and therefore could be expected to remain compatible with future Oxford Nanopore releases beyond the R10.4.1 model. We provide the raw POD5 signal data for the four benchmarking strains and the 31 diverse strains as a publicly available dataset for algorithm development for prokaryotic methylation. Future algorithm development in prokaryotic R-M could attempt to predict *de novo* motif specificity from methyltransferase gene sequence alone. In conclusion, MicrobeMod can be routinely used to rapidly describe prokaryotic methylation patterns and R-M systems in the nanopore sequencing of any prokaryotic isolate.

## Materials and Methods

### Nanopore sequencing

Genomic DNA from 35 microbial strains (**Table 1**) was extracted using the Qiagen DNeasy Blood and Tissue Kit following the recommended protocol with the following minor modifications: cultures were harvested at OD_600_ range of 1-1.5 before centrifugation, and the samples were incubated at room temperature for 4 minutes after adding 60 μL AE buffer for elution step. For a negative control we sequenced the NEB C2925H strain with the hsdR2 mutation which others have reported is completely methylation-deficient (Zymo). Library preparation for nanopore sequencing with the eluted gDNA was performed using Rapid Sequencing DNA V14 -Barcoding Kit (SQK-RBK114.24) from Oxford Nanopore Technologies following their official protocol. Twelve or fewer strains were loaded in each run with FLO-MIN114 R10.4.1 flow cells on a MinION Mk1C with a 72-hour run-limits, and sequencing runs were repeated for low-coverage samples after demultiplexing with MinKNOW on the Mk1C. Some samples were run with 4 kHz and some with 5 kHz frequencies after a switch by MinKNOW in mid-2023 (**Table 1**).

### Base calling and assembly

Base calling was performed on AWS p3.2xlarge instances with Dorado 0.4 (for 5 kHz run) and Dorado 0.3 (for 4 kHz runs), using a Docker image provided at the MicrobeMod GitHub.

For 5 kHz samples, the parameters used were:

~~~
dorado basecaller dna_r10.4.1_e8.2_400bps_sup@v4.2.0
--modified-bases-models
dna_r10.4.1_e8.2_400bps_sup@v4.2.0_5mC@v2,dna_r10.4.1_e8.2_400bps_sup
@v4.2.0_6mA@v3.
~~~

For 4 kHz samples, the parameters used were:

~~~
dorado basecaller res_dna_r10.4.1_e8.2_400bps_sup@v4.0.1 --emit-moves
--modified-bases-models
res_dna_r10.4.1_e8.2_400bps_sup@v4.0.1_5mC@v2,res_dna_r10.4.1_e8.2_40
0bps_sup@v4.0.1_6mA@v2.
~~~

Long-read genomes were assembled using Flye^47^ with the parameter: -g 5m –nano-hq –scaffold. The parameter --*asm*-*coverage 75* was passed for genomes with estimated depths of coverage greater than 150x. Assembled genomes were corrected with 2x150 bp Illumina reads using PolyPolish with default settings. Reads were mapped to the assembled reference genomes using the samtools fastq command (with parameter -T mv,MM,ML) to convert the BAM output of Dorado to FASTQ, and then minimap2^48^ was run with parameters --secondary=no -ax map-ont.

### Methylation motif-calling pipeline

The *call_methylation* workflow requires both a BAM file with R10.4.1+ nanopore reads (containing the MM,ML tags output that indicate methylation status as output by Dorado) mapped to a reference genome, and the FASTA file of corresponding reference genome. The first step calls Modkit pileup ([GitHub link](https://github.com/nanoporetech/modkit)), which is passed the optional “--methylation_confidence_threshold” parameter, with a default value of 0.66 in MicrobeMod, to generate a table of methylation frequencies per site. The resulting output table of per-site methylation frequencies is then filtered to sites above a minimum coverage per strand (“--min_coverage”, with a default of 10x) at least 66% of reads called as methylated (“--percent_methylation_cutoff”). Sites with at least 90% of reads called as methylated (“--percent_cutoff_streme”) are further prioritized for motif calling. 12 bp upstream and downstream of each methylated site are extracted as 24mers and passed to STREME^40^ for motif calling, identifying motifs statistically enriched compared to 100,000 randomly extracted 24mers from the same genome. STREME is then run with the parameter: --minw 4. MicrobeMod then filters motifs identified by STREME with an e-value cutoff of 0.1. Positions in each statistically significant motif with <80% of the sites corresponding to a single nucleotide are converted to ambiguous N characters, and trailing N’s are stripped from motifs. In our usage in this paper for low-coverage 10x-20x samples and/or subsamples, MicrobeMod was run with the min_coverage parameter modified from the default 10 to 3 (which corresponds to a stranded coverage of 3x for a total coverage of 6x, as methylation status is strand-specific).

### Restriction-modification annotation pipeline

The *annotate_rm* pipeline first calls genes and predicts proteins using prodigal^41^ for gene-calling and protein prediction (default settings). Next, a set of HMMs from the DefenseFinder^1^ package for the identification of Type I, Type II, Type III, and Type IV methyltransferases and restriction enzyme genes are run using HMMER hmmsearch^42^ on predicted proteins with the parameter –cut_ga. Further, the following HMMs from the PFAM database^49^ are also run to distinguish DNA methyltransferases from a variety of methyltransferases with other specificities involved in other cellular processes: PCMT, UPF0020, Ubie_methyltran, Methyltransf_3, MTS, PrmA, MTS_N, Methyltransf_15, Methyltrans_SAM, AviRa, Methyltransf_25, Methyltransf_31. The program cath-resolve-hits^50^ is then run to identify the best domain-specific HMM hits for each gene, and only genes which more closely matched the first set of DNA methyltransferase HMMs are retained. The closest REBASE homolog is identified for each RM gene using BLASTP ^51^ against a custom database of enzymes annotated by REBASE^31^, retaining hits with average amino acid percent identity >50% .

## Acknowledgements

We thank James Knight for assistance with deploying base calling on AWS Batch, and Elise Ledieu-Dherbécourt, Ariela Esmurria, and Wajd Alsharif for cultivation of the strains used in this study. We thank Tyler Barnum, Charlie Gilbert, Stephanie Brumwell, Paul Carini, Michael Molla, and the entire Cultivarium team for helpful discussion and feedback. Cultivarium acknowledges support from Schmidt Futures as a Convergent Research Focused Research Organization (FRO).

## Notes

https://github.com/cultivarium/MicrobeMod

### Competing Interest Statement

The authors have declared no competing interest.

